# HERV-K HML-2 transcription in diverse cancers is related with cancer stem cell and epithelial-mesenchymal transition markers

**DOI:** 10.1101/451997

**Authors:** Audrey T. Lin, Cindy G. Santander, Fabricia F. Nascimento, Emanuele Marchi, Timokratis Karamitros, Aris Katzourakis, Gkikas Magiorkinis

## Abstract

Endogenous retroviruses (ERVs) are remnants of ancient retroviral infections that make up 8% of the human genome. Although these elements are mostly fragmented and inactive, many proviruses belonging to the HERV-K (HML-2) family, the youngest lineage in the human genome, have intact open reading frames, some encoding for accessory genes called *np9* and *rec* that interact with oncogenic pathways. Many studies have established that ERVs are transiently expressed in both stem cells and cancer, resulting in aberrant self-renewal and uncontrolled proliferation. *np9* and *rec* expression are significantly correlated with a range of cancer stem cell (CSC) and epithelial to mesenchymal transition (EMT) biomarkers, including cellular receptors, transcription factors, and histone modifiers. Surprisingly, these ERV genes are negatively correlated with genes known to promote pluripotency in embryonic stem cell lines, such as Oct4. These results indicate that HERV-K (HML-2) is part of the transcriptional landscape responsible for cancer cells undergoing the phenotypic switch that characterises EMT. The discovery of *np9* and *rec*’s correlation with CSC and EMT biomarkers suggest a yet undescribed role affecting the transitional CSC-like state in EMT and the shift towards cancer malignancy.

**Importance:** In this study, we find that human endogenous retrovirus HERV-K (HML-2)-encoded genes *np9* and *rec* are correlated with the expression of many biomarkers associated with cancer stem cells (CSC) and epithelial-mesenchymal transition (EMT). There has been a significant effort to develop novel treatments targeting CSC and EMT-specific signalling pathways and cell surface markers. This research describes HERV-K (HML-2) as interacting or being part of the regulatory network that make up reversible cell state switching in EMT. Our findings suggest these specific HERVs may be good candidate biomarkers in identifying the transitional CSC-like states that are present during the progression of EMT and cancer metastasis.

## Introduction

Human endogenous retroviruses (HERVs) are remnants of ancient retroviral infections that propagated in the germlines of our distant mammalian ancestors and are present in multiple copies and lineages in their host genomes [1]. HERVs stopped proliferating in our ancestors’ genomes millions of years ago, with one exception, HERV-K (HML-2) (referred here as HK2), which continued to expand after the human-chimpanzee split [2]. The majority of HERVs are defective, due to accumulation of random mutations, selection pressure from host-derived restriction factors, or recombination, resulting in frameshifts, premature stop codons, and large deletions that remove the coding capacity of the provirus [1]. Many HERVs are also tightly regulated by epigenetic modification [3]. HK2 integrations, on the other hand, have frequently more intact open reading frames compared to older HERVs. Some HK2 proviruses have the capacity to produce the accessory proteins Np9 and Rec depending on the presence (type I proviruses encoding for *np9*) or absence (type 2 proviruses encoding for *rec*) of a deletion in the *env* gene [4], [5]. These proteins are known to be highly expressed in a number of cancers, germ cell tumour cell lines, and human embryonic stem cells [6]. HK2-encoded proteins have been shown to have potential oncogenic function: Np9 and Rec have been shown to interact with cellular pathways linked to cancer [7], [8], [9], including the activation of the oncogene *myc* [8], and the Notch signalling pathway [8]. Rec protein functions in regulating viral gene expression by transporting viral mRNA from the nucleus into the cytosol [10], [6], and its expression has been found to interfere with the development of germ cells and also cause carcinoma in mice [10]. In addition, Np9 protein is reported to bind and regulate ubiquitin ligase MDM2, resulting in the repression of p53 tumour suppressor [11]. Transcripts of *np9* and *rec* are found in human embryonic stem cells and human induced pluripotent stem cell lines and are associated with maintenance of pluripotency [12], [13], and recently it has been demonstrated that the activation and silencing of a specific HERV-K LTR LTR5HS is associated with reciprocal up-and downregulation of hundreds of human genes, and can occur over long genomic distances [14].

Within a tumour, there is a small subset of biologically distinct and rare “tumour-initiating” cells with attributes that parallel the functional properties of normal stem cells, referred to as cancer stem cells (CSCs). CSCs have been proposed to be the source of all malignant cells in a primary tumour, and their presence also gives rise to metastatic growth. It has also been suggested that the failure to eliminate CSCs results in relapse and a drug-resistant phenotype of the tumour regrowth, following chemotherapy-induced remission [15]. Because of the tumour-initiating ability of CSCs and the association of stem cell behaviours by normal and neoplastic cells during the activation of epithelial to mesenchymal transition (EMT) [15], there has been a significant effort to develop novel treatments targeting CSC and EMT-specific signalling pathways and cell surface markers.

Since HK2 is transcribed in many different cancers, during specific stages of early development, and is involved in the maintenance of pluripotency in embryonic stem cells [16], [6], HK2 overexpression in seemingly diverse cancer types might reflect the stem cell-like identity of the tumour. Whether the upregulation of HK2 contributes to tumorigenesis and cancer progression or is simply an unrelated side-effect of increased cellular dysfunction is not very well understood. To better address this problem, we characterise in this study a list of transcriptional interactions between HK2 and specific components of stem cell/EMT-associated pathways.

## Materials and Methods

### Data Analysed

The method described in [17] was used in capturing *np9* and *rec*-specific transcripts in RNA-seq data. Raw FASTQ files of Illumina HiSeq RNA-seq reads, originally sequenced from 305 samples derived from primary solid tumour tissues and blood-derived normal tissues from different individuals, were downloaded from Cancer Genomics Hub (data now available at Genomics Data Commons, https://gdc.cancer.gov). A total of 15 different cancer types were analysed. Cancer types reported to be associated with HK2 expression were selected for study: breast cancer, lymphoma, lung cancer (lung adenocarcinoma and lung squamous cell carcinoma), ovarian cancer, prostate cancer, skin melanoma, liver hepatocellular carcinoma, and testicular germ cell cancer. The other 6 cancer types were selected by tumour type: melanoma (uveal melanoma), adenocarcinomas (colon, pancreatic, and thyroid adenocarcinomas), carcinomas (cervical carcinoma, cholangiocarcinoma) (**Table 1**). Normal matched controls are not available for the majority of the tumour samples in TCGA, so a selection of random normal non-tumour tissues (cervix, bile duct, bowel, liver, lung, ovary, pancreas, prostate, testis, and thymus) from TCGA and Illumina Human Body Map 2.0 was used. The non-diseased samples used for the comparison group (referred in the next as “NT”) were taken from.

**Table 1.**
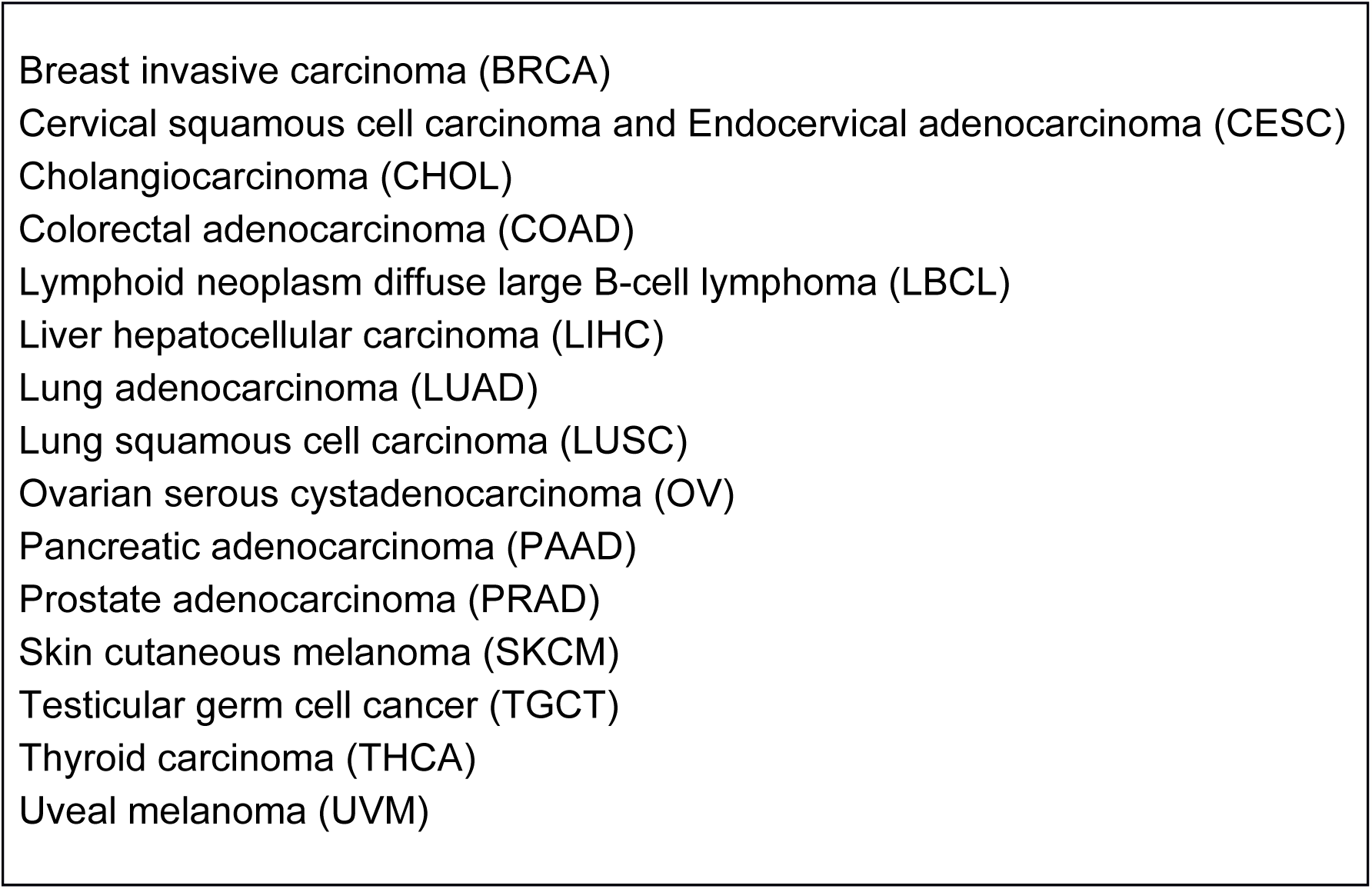
**Summary of The Cancer Genome Atlas (TCGA) RNA-seq data**

|  |
| --- |
| Breast invasive carcinoma (BRCA) |
| Cervical squamous cell carcinoma and Endocervical adenocarcinoma (CESC) |
| Cholangiocarcinoma (CHOL) |
| Colorectal adenocarcinoma (COAD) |
| Lymphoid neoplasm diffuse large B-cell lymphoma (LBCL) |
| Liver hepatocellular carcinoma (LIHC) |
| Lung adenocarcinoma (LUAD) |
| Lung squamous cell carcinoma (LUSC) |
| Ovarian serous cystadenocarcinoma (OV) |
| Pancreatic adenocarcinoma (PAAD) |
| Prostate adenocarcinoma (PRAD) |
| Skin cutaneous melanoma (SKCM) |
| Testicular germ cell cancer (TGCT) |
| Thyroid carcinoma (THCA) |
| Uveal melanoma (UVM) |

To analyse mRNA expression of EMT and CSC genes, the data was obtained from TCGA, provisional by using www.cbioportal.org.

### Testicular germ cell cancer as cancer stem cell reference

Testicular germ cell cancer is used as a CSC reference in this study because of the distinctly similar molecular and cellular phenotypes observed in germ cell tumours and human embryonic stem cells. These shared traits include lack of contact inhibition in culture, a rapid proliferation rate, high activity of telomerase, and similarities in gene expression and epigenetic patterns, like the upregulation of stem cell markers *P0U5F1* (Oct4), *NANOG*, and *S0X2*, and oncogenes *MYC* and *KLF4*, and the downregulation of the tumour suppressor *TP53* (reviewed in [10]).

### Analysis of EMT and CSC genes from TCGA datasets

For expression analysis of EMT and CSC genes, the data was obtained from TCGA by using www.cbioportal.org. EMT and CSC genes analysed were classified depending on either being reportedly overexpressed (or upregulated) and transcriptionally repressed (downregulated) in stem cells and EMT (**Table 2, Supplementary Information**). The gene groups are classified as follows: EMT_up genes (77 genes); EMT_down genes (13 genes); CSC_up genes (47 genes); CSC_down genes (8 genes).

**Table 2.** **Lists of gene groups analysed**

| Group | Genes |
| --- | --- |
| **EMT – upregulated genes (EMT_up)** | AKT1, ALDH1A1, AXIN2, BMI1, BMP4, CD44, CSN2, CTNNB1, EHMT2, EZH2, FGF2, FOXC2, GLI1, GRHL2, GSC, GSK3B, HDAC1, HDAC2, HDAC3, HIF1A, IL6, JAK1, JAK2, KAT5, KDM1A, KDM5B, KLF4, KLF8, LATS2, LBX1, LIN28A, LOXL2, MAPK1, MYC, NANOG, NODAL, NOG, NOTCH1, NOTCH2, NOTCH3, NOTCH4, OVOL1, OVOL2, PAK1, PDGFRB, POU5F1, PRDM14, PRRX1, QKI, SALL4, SETDB1, SHH, SIN3A, SIRT1, SIX1, SMAD2, SMAD3, SMAD4, SMO, SNAI1, SNAI2, SOX2, SRSF2, |
|  | STAT3, SUV39H1, SUZ12, T, TCF3, TCF4, TGFB1, TGFB2, TWIST1, WNT2, YBX1, ZEB1, ZEB2, ZFP42 |
| **CSC – upregulated genes (CSC_up)** | ALDH1A1, APC, AXIN2, CD34, CD44, CTNNB1, DLL4, EP300, FGF2, KDM1A, KDM5B, KLF2, KLF4, KLF8, LATS2, LIN28A, MAPK1, MYC, NANOG, NODAL, NOG, NOTCH1, NOTCH2, NOTCH3, NOTCH4, NRB01, POU5F1, PROM1, SALL4, SHH, SIN3A, SMAD2, SMAD3, SMAD4, SMO, SOX18, SOX2, STAT3, SUV39H1, TCF3, TCF4, TGFB1, TGFB2, THY1, WNT2, YBX1, ZFP42 |
| **EMT – downregulated genes (EMT_down)** | APC, CD24, CD38, CDH1, CTNND1, ENAH, ESRP1, ESRP2, FBXL14, HSP90AA1, MDM2, PRKD1, TP53 |
| **CSC – downregulated genes (CSC_down)** | BMP4, CD24, CD38, CTNND1, GSK3B, IL6, SETDB1, SUZ12 |

**Note:** EMT – epithelial to mesenchymal transition, CSC – cancer stem cells.
Upregulated genes – genes in this group are reported by the literature to be upregulated in the context of EMT/CSC. Downregulated genes – genes in this group are reported by the literature to be downregulated in the contexts of EMT/CSC.

### Calculation of transcriptional distance

Testicular germ cell cancer (TGCT) is used as a CSC reference because of distinctly similar molecular and cellular phenotypes observed in germ cell tumours and human embryonic stem cells [18]. For the TCGA-TGCT RNA-seq dataset, the mean expression value was calculated for each gene, and the similarity of the expression profile of each tumour sample to the expression profile of the CSC was calculated using Formulas 1 and 2. In Formula 1, within each tumour sample per cancer, φ is the distance from the cancer stem cell (CSC) phenotype (either EMT or stem cell genes) in EMT/CSC_up genes in a given tumour sample, μ*g*_*tc*_ is the average expression of the gene in all the TCGA-TGCT samples, and *g*_*z*_ is the expression of the gene in a given tumour sample. Therefore, the more negative φ is, the more similar the expression profile of the tumour sample is to the CSC phenotype. In either EMT_down or CSC_down genes, Formula 2 is used, where ψ is the distance from the CSC phenotype (either EMT or stem cell genes), and the more positive ψ is, the more similar the expression profile of the given tumour sample to the CSC phenotype.

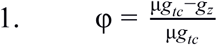

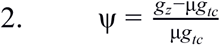

Because some of the transcript counts were zero or negligible, transformation of these values were necessary in order to conduct further statistical analyses to find the strength of association between HK2 expression in each tumour sample to the EMT and CSC gene expression profiles (represented by φ and ψ ). These values were transformed using formulas 3 and 4, respectively. The integers 10,000+1 used in these formulas is to account for the non-negative values that would otherwise result with the log transformation.

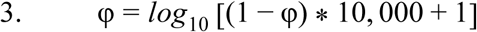

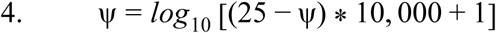

### Statistical Analysis

The Wilcoxon signed rank-sum test was used to compare the relative expression of the HK2 genes from each of the unmatched samples within each cancer type compared to NT. Multiple linear regression with interaction terms was used to estimate the association between the expression of the HK2 targets and up (φ) and down (ψ) EMT and CSC genes in the different cancers. The chi-square test of independence was performed to determine the statistical significance of the correlations between the expression of *np9* or *rec* and activity of functional categories of CSC/EMT genes. The gene categories consist of cell surface receptors (10 genes), transcription factors (29 genes), and genes categorised as “Other”: negative regulators (3 genes), histone modifiers (11 genes), kinases (5 genes), metabolic enzyme (1 gene), ubiquitin ligase (1 gene), splicing factors (2 genes), receptor ligands (4 genes), and cell adhesion molecules (3 genes). For all statistical methods, associations with p-value < 0.05 were considered statistically significant. All statistical analyses were performed in R version 3.3.1 [19].

## Results

### HK2 expression is high in cells with CSC/EMT-like transcriptional profiles

HK2 is generally overexpressed across different cancer types[17], thus we wanted to determine whether there is a correlation between *np9* and *rec* activity and the expression of genes associated with CSCs and EMT. High transcription of *np9* and *rec* is only observed in cancers with highly similar EMT/CSC transcriptional profiles (values closer to (1.6, *y*)), and a similar pattern is also seen in *pol* and HERVK-113 LTRs (**Fig. 1**). These results suggest that the CSC-like transcriptional profile is not sufficient by itself as an explanation for high HK2 transcription. The specific genes that could be independently associated with the observed transcriptional pattern between CSC/EMT genes and HK2 were next investigated.

**Figure 1.**
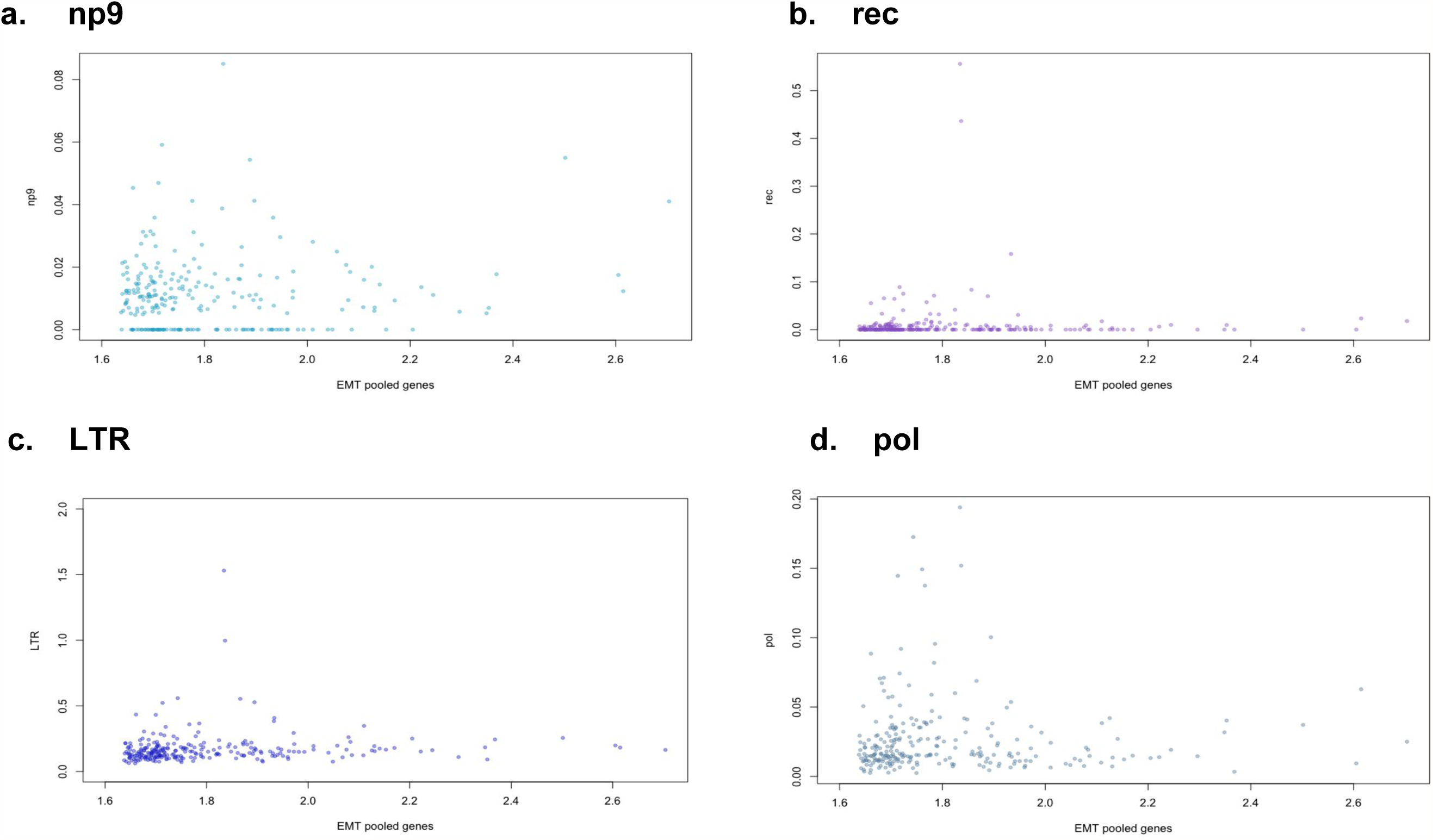
*np9* and *rec* expressed in cancers that have highly similar epithelial-mesenchymal transition (EMT) and cancer stem cell (CSC) gene expression signature, x-axis indicates the pooled gene expression signature of all the EMT/CSC genes within all the tumour samples. Values closer to (1.6, *y*) indicates more similarity to the EMT/CSC phenotype. y-axis indicates the relative expression of **(a.)** *np9*; **(b.)** *rec*; **(c.)** HERV-K113 LTR; **(d.)** *pol* in all tumour samples – these values have been transformed by square root.

**Figure 2.**
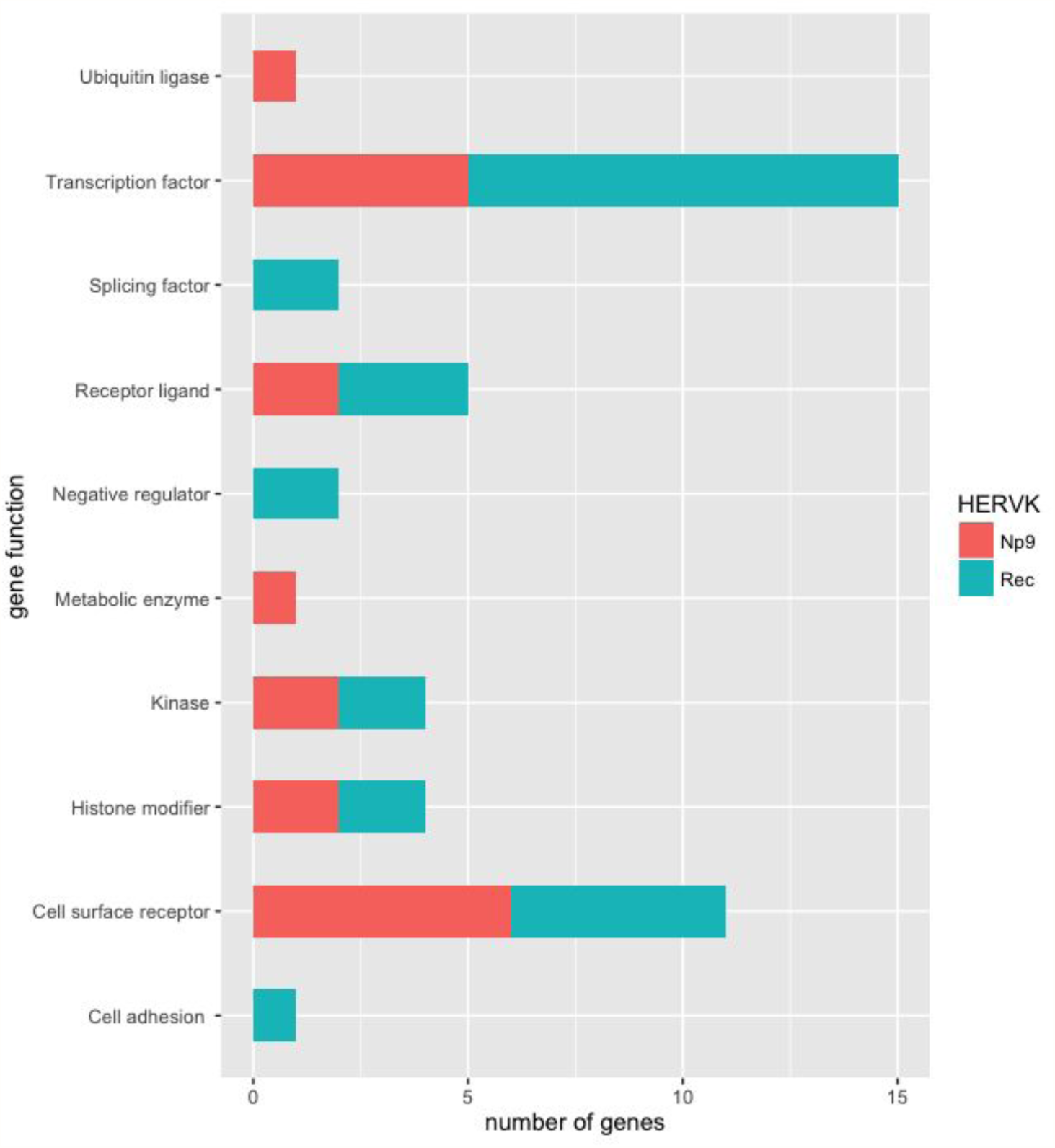
Breakdown of different gene categories. x-axis indicates the number of genes *np9* and *rec* are significantly correlated with; y-axis indicates the generalised function of the host genes that are significantly correlated with *np9* and *rec* expression.

### Independent correlates of HK2 expression and CSC/EMT gene expression

The overall transcriptional landscape was described for the CSCs and the CSC-like transitory state in EMT. The results indicate that HK2 transcription is significantly correlated with a number of specific CSC/EMT genes: there are 22 significant positive and negative correlations between the expression of *np9* and EMT/CSC genes, and 30 significant correlations between the expression of *rec* and the EMT/CSC genes (**Table 3**).

**Table 3.** (a) *np9* is significantly correlated with specific CSC/EMT upregulated genes

| Gene function category | Gene | Group | Coefficient | p-value |
| --- | --- | --- | --- | --- |
| Transcription factor | NODAL | EMT | 0.13 | $2.25 \times 10^{-3}$ |
| | OVOL2 | EMT | -0.05 | $1.74 \times 10^{-3}$ |
| | T (Brachyury) | EMT | 0.07 | $9.23 \times 10^{-3}$ |
| | CTNNB1 (β-catenin) | CSC | 0.26 | $1.35 \times 10^{-2}$ |
| | TCF3 (E47) | EMT | -0.52 | $6.89 \times 10^{-7}$ |
| | TCF3 (E47) | CSC | 0.26 | $1.35 \times 10^{-2}$ |
| Cell surface receptor | NOTCH4 | CSC | 0.42 | $1.50 \times 10^{-4}$ |
| | CD133 (PROM1) | CSC | 0.06 | $5.00 \times 10^{-3}$ |
| | NOTCH1 | CSC | -0.2 | $2.85 \times 10^{-2}$ |
| | NOTCH2 | CSC | -0.24 | $2.00 \times 10^{-2}$ |
| | CD44 | CSC | -0.11 | $2.49 \times 10^{-2}$ |
| Receptor ligand | DLL4 | CSC | -0.31 | $4.20 \times 10^{-4}$ |
| | WNT2 | CSC | -0.08 | $3.67 \times 10^{-4}$ |
| Histone modifier | EHMT2 (G9a) | EMT | 0.4 | $9.47 \times 10^{-4}$ |
| | SUV39H1 | CSC | 0.47 | $5.81 \times 10^{-5}$ |
| Kinase | EP300 (p300) | CSC | -0.3 | $3.52 \times 10^{-2}$ |
| | LATS2 | CSC | 0.21 | $2.83 \times 10^{-2}$ |
| Metabolic enzyme | ALDH1A1 | EMT | -0.08 | $2.54 \times 10^{-2}$ |

**(b) *rec* is significantly correlated with specific CSC/EMT upregulated genes**
| Gene function category | Gene | Group | Coefficient | p-value |
| --- | --- | --- | --- | --- |
| Transcription factor | GRHL2 | EMT | -0.04 | $2.30 \times 10^{-2}$ |
| | GSC (Goosecoid) | EMT | -0.08 | $1.61 \times 10^{-4}$ |
| | MYC | CSC | 0.15 | $2.49 \times 10^{-2}$ |
| | POU5F1 (Oct4) | CSC | -0.12 | $1.46 \times 10^{-2}$ |
| | POU5F1 (Oct4) | EMT | -0.12 | $1.69 \times 10^{-2}$ |
| | SMAD2 | EMT | -0.37 | $1.07 \times 10^{-2}$ |
| | STAT3 | EMT | -0.35 | $2.40 \times 10^{-2}$ |
| | TCF4 (E2-2) | CSC | -0.2 | $9.65 \times 10^{-3}$ |
| | TWIST1 | EMT | -0.12 | $5.98 \times 10^{-3}$ |
| | CTNNB1 (β-catenin) | EMT | 0.66 | $2.53 \times 10^{-5}$ |
| | TCF3 | EMT | -0.31 | $1.17 \times 10^{-2}$ |
| Cell surface receptor | PDGFRB | EMT | 0.2 | $1.36 \times 10^{-2}$ |
| | CD44 | EMT | 0.28 | $1.18 \times 10^{-7}$ |
| | CD44 | CSC | 0.21 | $4.50 \times 10^{-5}$ |
| | NOTCH4 | CSC | 0.27 | $3.08 \times 10^{-2}$ |
| | SMO | EMT | 0.15 | $1.18 \times 10^{-2}$ |
| Receptor ligand | WNT2 | CSC | -0.07 | $5.09 \times 10^{-3}$ |
| | FGF2 | CSC | -0.11 | $3.46 \times 10^{-2}$ |
| | BMP4 | EMT | -0.1 | $4.32 \times 10^{-2}$ |
| Histone modifier | SUV39H1 | EMT | -0.41 | $5.85 \times 10^{-4}$ |
| Kinase | LATS2 | CSC | 0.27 | $9.97 \times 10^{-3}$ |
| Negative regulator | CSN2 | EMT | 0.03 | $1.93 \times 10^{-2}$ |

**(c) *np9* is significantly correlated with specific CSC/EMT downregulated genes**
| Gene function category | Gene | Group | Coefficient | p-value |
| --- | --- | --- | --- | --- |
| Cell surface receptor | CD38 | CSC | 12.8 | $7.97 \times 10^{-4}$ |
| | CD38 | EMT | 5.14 | $2.51 \times 10^{-3}$ |
| Receptor ligand | BMP4 | CSC | 2 | $4.96 \times 10^{-2}$ |
| Ubiquitin ligase | FBXL14 | EMT | 11.3 | $1.45 \times 10^{-2}$ |

**(d) *rec* is significantly correlated with specific CSC/EMT downregulated genes**
| Gene function category | Gene | Group | Coefficient | p-value |
| --- | --- | --- | --- | --- |
| Cell surface receptor | CD24 | CSC | 2.4 | $6.90 \times 10^{-3}$ |
| Histone modifier | SUZ12 | CSC | 43.2 | $1.92 \times 10^{-4}$ |
| Kinase | PRKD1 | EMT | -0.33 | $4.40 \times 10^{-2}$ |
| Negative regulator | APC | EMT | 7.37 | $4.21 \times 10^{-2}$ |
| Splicing factor | ESRP2 | EMT | 4.2 | $5.82 \times 10^{-6}$ |
| | ESRP1 | EMT | 14.7 | $3.01 \times 10^{-3}$ |
| Cell adhesion | CTNND1 | EMT | -8.67 | $6.26 \times 10^{-4}$ |

Summary of significant correlations between upregulated genes within the CSC or EMT groups with **(a.)** *np9* and **(b.)** *rec*. Summary of significant correlation between downregulated genes within the CSC or EMT groups with **(c.)** *np9* and **(d.)** *rec*. Parentheses () indicates other names for the same gene.

*np9* transcription is significantly correlated with the transcription of 5 transcription factor (NODAL, OVOL2, TBXT (Brachyury), CTNNB1 (β-catenin), and TCF3), 6 cell surface receptors (CD38, CD133, NOTCH1, NOTCH2, CD44, NOTCH4), 3 receptor ligands (DLL4, BMP4, WNT2), 2 histone modifiers (EHMT2 (G9a), SUV39H1), 2 kinases (EP300, LATS2), 1 metabolic enzyme (ALDH1A1), and 1 ubiquitin ligase (FBXL14) (**Table 3a, Table 3c, Fig. 3**). *rec* transcription is significantly and independently correlated with 10 transcription factors (GRHL2, GSC (Goosecoid), MYC, POU5F1 (Oct4), SMAD2, STAT3, TCF4, TWIST1, CTNNB1 (β-catenin), and TCF3), 5 cell surface receptors (CD24, PDGFRB, CD44, NOTCH4, and SMO), 3 receptor ligands (FGF2, BMP4, and WNT2), 2 histone modifiers (SUZ12 and SUV39H1), 2 kinases (PRKD1 and LATS2), 2 negative regulators (APC and CSN2), 2 splicing factors (ESPR1/2), and 1 cell adhesion molecule (CTNND1) (**Table 3b, Table 3d, Fig. 3**). HK2 transcription is most prominently correlated with the expression of genes encoding for transcription factors and cell surface receptors, (**Fig. 3**). However, only *rec* transcription is independently and significantly correlated with expression of gene types that fall into the functional categories of cell surface receptors, transcription factors, and “Other” gene types (**Table 4a**), while *np9* is not independently correlated with any of these functional categories (**Table 4b**).

**Figure 3.**
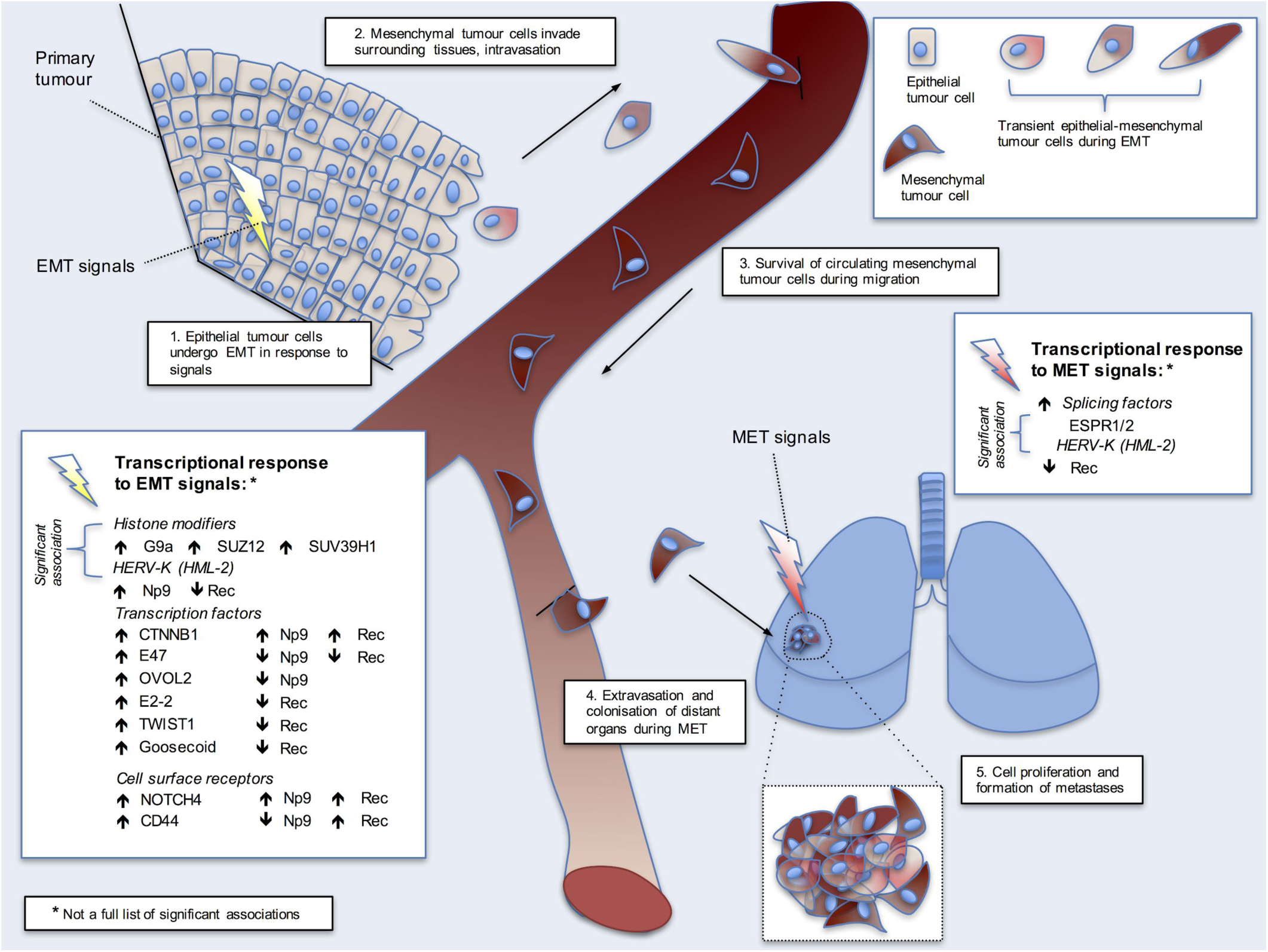
Model of *np9* and *rec* transcriptional activity during epithelial-mesenchymal transition (EMT). 1. Epithelial tumour cells undergo EMT in response to signals that include the activity of transcriptional response of histone modifiers, transcription factors, and cell surface receptors. When the histone modifiers G9a, SUZ12, and SUV39H1 are upregulated, there is an upregulation of *np9*, and a downregulation of *rec*. The inverse is seen with the upregulation of cell surface receptor CD44 with a downregulation of *np9* and an upregulation of *rec*. When the cell surface receptor gene NOTCH4 is upregulated, there is also an upregulation of *np9* and *rec*. The upregulation of transcription factor CTNNB1 occurs with the co-expression of *np9* and *rec*; the upregulation of E47 occurs with the co-repression of *np9* and *rec*; the upregulation of OVOL2 occurs with the downregulation of *np9*; and the upregulation of E2-2, TWIST1, and Goosecoid all result with the downregulation of *rec*. 2. Mesenchymal tumour cells then invade the surrounding tissues, and intravasation occurs in vasculature. 3. The circulating mesenchymal tumour cells survive as they migrate within vasculature. 4. Extravasation of the mesenchymal tumour cells and colonisation of a new site. These tumour cells undergo mesenchymal-epithelial transition (MET) in response to MET-specific signals that include the transcriptional response of splicing factors, specifically the upregulation of splicing factors ESPR1/2 and the downregulation of *rec*. 5. Cellular proliferation and the formation of metastases at the new site.

**Table 4.** Chi-square contingency tables showing significant correlations between different functional gene categories and transcription of **(a.)** *rec* and **(b.)** *np9*.

| Contingency table of significant correlation between Rec and gene categories |  |  |  |
| --- | --- | --- | --- |
| Gene Category | Positive | Negative | Not significant |
| **Cell surface receptor** | 4 (1.14) [7.14] | 1 (2.71) [1.08] | 5 (6.14) [0.21] |
| **Transcription factor** | 1 (3.31) [1.62] | 8 (7.87) [0.0] | 20 (17.81) [0.27] |
| **Other** | 3 (3.54) [0.08] | 10 (8.41) [0.30] | 18 (19.04) [0.06] |
| **Column Total** | 8 | 19 | 43 |
| $X^2 = 10.76^{**}$ | | | |
| $**p = 0.029$ | | | |

| Contingency table of significant correlation between *np9* and gene categories |  |  |  |
| --- | --- | --- | --- |
| Gene Category | Positive | Negative | Not significant |
| **Cell surface receptor** | 1 (1.03) [0.0] | 4 (1.41) [4.73] | 4 (6.56) [1.0] |
| **Transcription factor** | 2 (3.43) [0.60] | 2 (4.71) [1.56] | 26 (21.86) [0.79] |
| **Other** | 5 (3.54) [0.60] | 5 (4.87) [0.0] | 21 (22.59) [0.11] |
| **Column Totals** | 8 | 11 | 51 |
| $\chi^2 = 9.38^{**}$<br>$^{**}p = 0.052$ | | | |

The gene categories “cell surface receptor” contains 10 genes and “transcription factor” contains 29 genes. The gene category “Other” consists of the following gene categories: negative regulator (3 genes), histone modifier (11 genes), kinase (5 genes), metabolic enzyme (1 gene), ubiquitin ligase (1 gene), splicing factor (2 genes), receptor ligand (4 genes), cell adhesion (3 genes). Each cell contains the following information: the observed cell totals, the expected cell totals in parentheses (), and the chi-square (X^2^) statistic in brackets [ ].

Chi-square contingency tables showing significant correlations between different gene categories and transcription of *np9*. The gene categories “cell surface receptor” contains 9 genes and “transcription factor” contains 30 genes.

The gene category “Other” consists of the following gene categories: negative regulator (3 genes), histone modifier (11 genes), kinase (5 genes), metabolic enzyme (1 gene), ubiquitin ligase (1 gene), splicing factor (2 genes), receptor ligand (4 genes), cell adhesion (3 genes). Each cell contains the following information: the observed cell totals, the expected cell totals in parentheses (), and the chi-square (X^2^) statistic in brackets [ ]

The significant correlation between *rec* activity with overexpression of the cell surface molecules CD44 (EMT group: p = 1.18 x 10^−7^; CSC group: p = 4.50 x 10^−5^) and CD24 underexpression (CSC group: p = 6.90 x 10^−3^) indicates an association with the CD44^high^/CD24^low^ phenotype, known to be enriched in breast cancer CSCs [20], [21]. Strikingly, there is conflicting expression states between *np9* and *rec* and CD44 expression, where *rec* is positively correlated (CSC group: p = 4.50 x 10^−5^), and *np9* is negatively correlated (CSC group: p = 2.49 x 10^−2^). Summaries of all the significant correlations between upregulated and downregulated genes within the CSC/EMT groups with *np9* and *rec* expression are in **Table 3**.

## Discussion

Previous studies have also shown that HK2 expression is high in specific stages of early embryonic development [6], [12], [13], [17]. HK2 is upregulated in tumours with greater transcriptional similarity with testicular germ cell cancer, a type of malignancy known for a strong stem-cell identity. This suggests that overexpression of HK2 can be found in diverse cancers that have a stem cell-like transcriptome. Together these observations indicate that HK2 transcription is a strong marker of CSC identity within a tumour. This is the first study, to our knowledge, that associates *np9* and *rec* activity with the expression of many human-specific transcriptional regulatory genes and signalling pathway genes implicated in the maintenance of cellular self-renewal and cancer malignancy through the cell state changes of EMT.

The transient expression of HK2 may be part of the transcriptional repertoire that shapes stem cell behaviour in CSCs. For example, *rec* is associated with the CD44^high^CD24^low^ antigenic profile in the cancer types surveyed. This phenotype is linked with increased malignancy and the presence of CSCs in breast tissues [20], [21], [22]. CD44^high^CD24^low^ cells also share many features with normal stem cells and are shown to be highly tumourigenic when injected into immunocompromised mice [22]. In addition, *rec* correlates with EMT biomarkers controlled by the ESPR (epithelial splicing regulatory protein) proteins, which regulates the splicing of cell-specific mRNAs in epithelial cells [23] and when highly expressed, promotes an epithelial phenotype. ESPR1 and its paralog ESPR2 differentially splice the transcripts of CTNND1 (catenin delta-1, also known as p120 catenin), CD44, and ENAH into epithelium-specific shorter isoforms. The absence of ESPR1/2 results in the longer isoforms being transcribed, resulting in a mesenchymal cell phenotype. CTNND1 functions as a cell adhesion and signal transduction molecule and the longer, epithelial-specific isoform of CTNND1 stabilises E-cadherin and promotes cell-cell adhesion [23]. Negative correlations were observed between *rec* expression and both ESPR1 and ESPR2, although out of the genes that ESPR1/2 regulates, there is a negative association between transcription of *rec* and CTNND1, and a positive association with CD44. The results of this study suggest that the cells where *rec* is transcriptionally active also repress the transcription of ESPR1/2, perhaps due to the tumour cells favouring the mesenchymal cell state and stem cell markers. It may be useful in future studies to shed more light on HK2 transcriptional presence within the more epithelial or mesenchymal phenotype by determining the isoform type of CTNND1 and CD44 that is transcribed in concert with high *rec* activity.

There was also a significant association between HK2 transcription and the deregulation of members of the Notch family of receptor proteins. In mammals, one of the five known ligands that bind to Notch includes DLL4, which interacts specifically with NOTCH1 or NOTCH4, activating signal transduction and promoting vascular development and self-renewal in stem cells [24]. Interestingly, there is an inverse relationship between expression of *np9* and DLL4 in CSC genes, yet there is a positive correlation between NOTCH4 and *np9*. In addition, the *np9*-NOTCH4 upregulation is also positively correlated with the expression of growth differentiation factor NODAL, which has been associated with a more aggressive melanoma phenotype [24]. There is also some evidence that breast CSCs are more resistant to chemotherapy compared to more differentiated cancer cells [25], and this could be partly explained by preferential NOTCH4 activation over NOTCH1 signalling in breast CSC populations compared to differentiated cells [26]. It would be interesting to explore in future studies whether Np9 protein physically interacts with Notch4 protein or its effectors, or impacts their transcription in *trans*. Although there was no significant correlation observed between *np9* and the other Notch receptors, it has been reported that in leukaemia progenitor/stem cell lines, Np9 protein interacts with NOTCH1 and other important signalling genes implicated in cancers, namely AKT1, MYC, MAPK1, and CTNNB1 [8].

A surprising result of this study is the negative correlation between *rec*, *pol*, and POU5F1 (Oct4) (**Table 3b, Table S1b**), a transcription factor necessary for self-renewal of undifferentiated stem cells and cellular reprogramming of differentiated cells to a pluripotent state [27], [28]. It has been previously reported that hypomethylation of HK2 LTRs and the transactivation of Oct4 synergistically promotes expression of HK2 [6], however this does not seem to be the case for cancer tumours. *np9* is positively correlated with EHMT2 (G9a), a histone-lysine N-methyltransferase that regulates cellular self-renewal in early development by silencing the expression of Oct4, NANOG, and DNMT3L [28], [29]. In addition, G9a also has the ability to silence the transcription of tumour suppressor genes that would consequently induce EMT. The deactivation of Oct4 through heterochromatinisation appears as a complex, multistep process involving the activity of numerous transcriptional repressors, including G9a which functions as both a methylase and also a recruiter of histone deacetylases and other histone methylases to the Oct4 promoter [28]. It is likely that HK2 expression in the maintenance of self-renewal is more complex than previously thought.

The results of this study indicate that there is not a direct relationship between known “core” pluripotency factors and HK2. Instead, this relationship is likely to be a far more complex interactive process, which might depend on the sensitivity of CSCs and tumour cells to extrinsic factors within the dynamic physical tissue microenvironment, as well as with stochastic variation in gene expression levels that promote cell state changes [30]. Within a heterogeneous tumour population, cells with stem cell-like properties and metastatic potential may exist in a spectrum. For example, it has been suggested that CSCs are not distinct entities but rather malignant cells that transiently acquire stem cell-like characteristics as a result of EMT [30]. In EMT, epithelial cells gain and disseminate new invasive behaviours such as motility, environmental adaptability, and self-renewal that are not only associated with metastatic growth but also with the stem cell phenotype. EMT appears to be a dynamic state, with “metastable” hybrid cells expressing both mesenchymal and epithelial traits, with the ability to induce or reverse the process [31], [32]. This intermediate, plastic, “partial EMT” state observed in fibrosis, development, and cancer is initiated and maintained by transcriptional regulation, which is possibly influenced by aberrant expression of previously silenced HK2 insertions and other transposable elements that promote the misregulation of host genes [33]. The activation of HK2 activity could be an advantageous trait selected for during the tumour’s evolution, because the derepression of *np9* and *rec*’s oncogenic potential would favour the tumour’s growth.

A fundamental property in early development and normal cell and tissue differentiation is reversible cell state switching, and the transient microenvironments generated are ordered interactions between thousands of genes in each cell, all within an intricate gene regulatory network [30] that HK2 is a part of (**Fig. 4**) [14]. HK2, and specifically its genes *np9* and *rec*, would potentially be good candidate biomarkers in identifying the transitional cancer stem cell-like state in EMT and the shift towards cancer malignancy.

**Figure 4.**
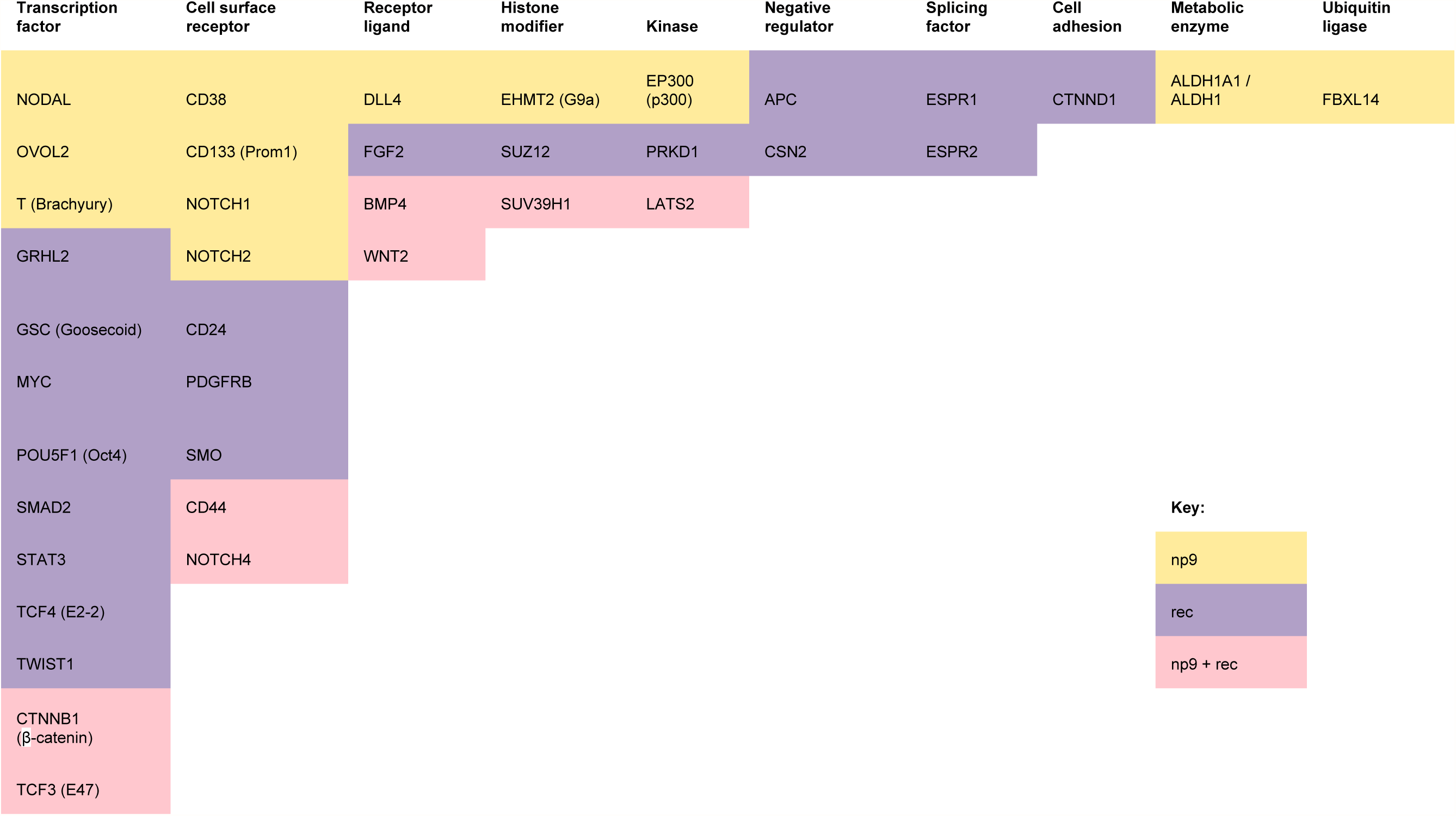
Host gene function categories significantly correlated with HK2 gene expression. Genes that are significantly correlated with the expression of *np9* and/or *rec* are categorised according to different host functions. The host genes that only *np9* is significantly correlated with are in the yellow cells, the host genes that only *rec* is significantly correlated with are in the purple cells, and the genes that both *np9* and *rec* are significantly correlated with are in the pink cells.

Limitations: The method for quantifying HK2 transcripts [17] is under validation and the results should be considered as preliminary/pending verification from a validation process.

## Acknowledgements

Thank you to Robert Belshaw, Tim Coulson, Samuel Campbell, and the Palaeovirology group for helpful discussions and comments.

## Funding Statement

GM and TK have been supported by an MRC fellowship to GM (MR/K010565/1); CS is funded by Comisión Nacional de Investigación Científica y Tecnológica, Gobierno de Chile; FFN is funded by The Royal Society and British Academy - Newton International Fellowship (140338); and AK is funded by The Royal Society. The funders had no role in study design, data collection and analysis, or preparation of the manuscript.

## Ethics Approval and Consent to Participate

All human data used were obtained through The Cancer Genome Atlas and used according to the Data Use Certification as stipulated by the TCGA Data Access Committee and dbGaP authorisation (https://dbgap.ncbi.nlm.nih.gov/aa/). Permission to use controlled-access data was approved by dbGaP under project #7621.

